# Structure of scavenger receptor SCARF1 and its interaction with lipoproteins

**DOI:** 10.1101/2023.11.08.566208

**Authors:** Yuanyuan Wang, Fan Xu, Guangyi Li, Chen Cheng, Bowen Yu, Ze Zhang, Dandan Kong, Fabao Chen, Yali Liu, Zhen Fang, Longxing Cao, Yang Yu, Yijun Gu, Yongning He

## Abstract

SCARF1 (Scavenger receptor class F member 1, SREC-1 or SR-F1) is a type I transmembrane protein that recognizes multiple endogenous and exogenous ligands such as modified low-density lipoproteins (LDL) and is important for maintaining homeostasis and immunity. But the structural information and the mechanisms of ligand recognition of SCARF1 are largely unavailable. Here we solve the crystal structures of the N-terminal fragments of human SCARF1, which show that SCARF1 forms homodimers and its epidermal growth factor (EGF)-like domains adopt a long-curved conformation. Then we examine the interactions of SCARF1 with lipoproteins and are able to identify a region on SCARF1 for recognizing modified LDLs. The mutagenesis data show that the positively charged residues in the region are crucial for the interaction of SCARF1 with modified LDLs, which is confirmed by making chimeric molecules of SCARF1 and SCARF2. In addition, teichoic acids, a cell wall polymer expressed on the surface of gram-positive bacteria, are able to inhibit the interactions of modified LDLs with SCARF1, suggesting the ligand binding sites of SCARF1 might be shared for some of its scavenging targets. Overall, these results provide mechanistic insights into SCARF1 and its interactions with the ligands, which are important for understanding its physiological roles in homeostasis and the related diseases.

## Introduction

Scavenger receptor (SR) was first discovered in late 1970s during the studies regarding the accumulation of low-density lipoprotein (LDL) in macrophages in atherosclerotic plaques of patients who lack LDL receptors [1, 2]. Up to date, SR family includes a large number of cell surface proteins that can be classified into more than ten classes (class A-L) based on the structural similarities [3]. SRs bind a wide range of endogenous and exogenous ligands including modified lipoproteins, damaged or apoptotic cells and pathogenic microorganisms [4–8] and play important roles in maintaining homeostasis, host defense and immunity, and have been linked to diseases such as cardiovascular diseases, Alzheimer’s disease and cancer [9–13].

Scavenger receptor class F (SR-F) has two known members, SCARF1 and SCARF2 [3] (Fig. 1A). SCARF1 was identified in cDNA libraries from human umbilical vein endothelial cells as a receptor for modified LDLs, thus also named SREC-1 (Scavenger Receptors expressed in Endothelial Cells) [14]. It is expressed on the surface of endothelial cells, macrophages, and dendritic cells and distributed in organs such as heart, liver, kidney and spleen [15]. SCARF1 recognizes modified LDLs, including acetylated LDL (AcLDL), oxidized LDL (OxLDL) and carbamylated LDL [14, 16, 17]. Previous studies have shown that the SCARF1-mediated degradation of AcLDL accounts for 60% of the amounts of AcLDL degraded by the pathway independent of scavenger receptor class A (SR-A), suggesting that SCARF1 may play a key role in the development of atherosclerosis in concert with SR-A in some situations [15]. SCARF1 expressed on antigen-presenting cells such as dendritic cells can recognize and internalize heat shock protein-bound antigens and activate adaptive immune responses [18–20]. It may also cooperate with the Toll-like receptors to mediate the cytokine production [7, 21, 22]. SCARF1-knockout mice can develop symptoms similar to systemic lupus erythematosus disease and lead to accumulation of apoptotic cells in the immune organs, suggesting that it is involved in the removal of apoptotic cells and maintaining homeostasis [23]. SCARF1 may also associate with the extravasation of leukocytes from circulation into inflamed tissues during injury or infection, thus having a role in the inflammatory changes in vessel walls and the initiation of atherosclerosis [24, 25]. Recent data also show that SCARF1 is down-regulated in hepatocellular carcinoma and loss of SCARF1 is associated with poorly differentiated tumors [26]. SCARF2 has 35% sequence identity and similar organ distribution with SCARF1 [24]. Genetic analysis suggests that SCARF2 is linked to a rare disease called van den Ende-Gupta syndrome [27, 28], which may provide clues for the physiological roles of this molecule.

**Figure 1.**
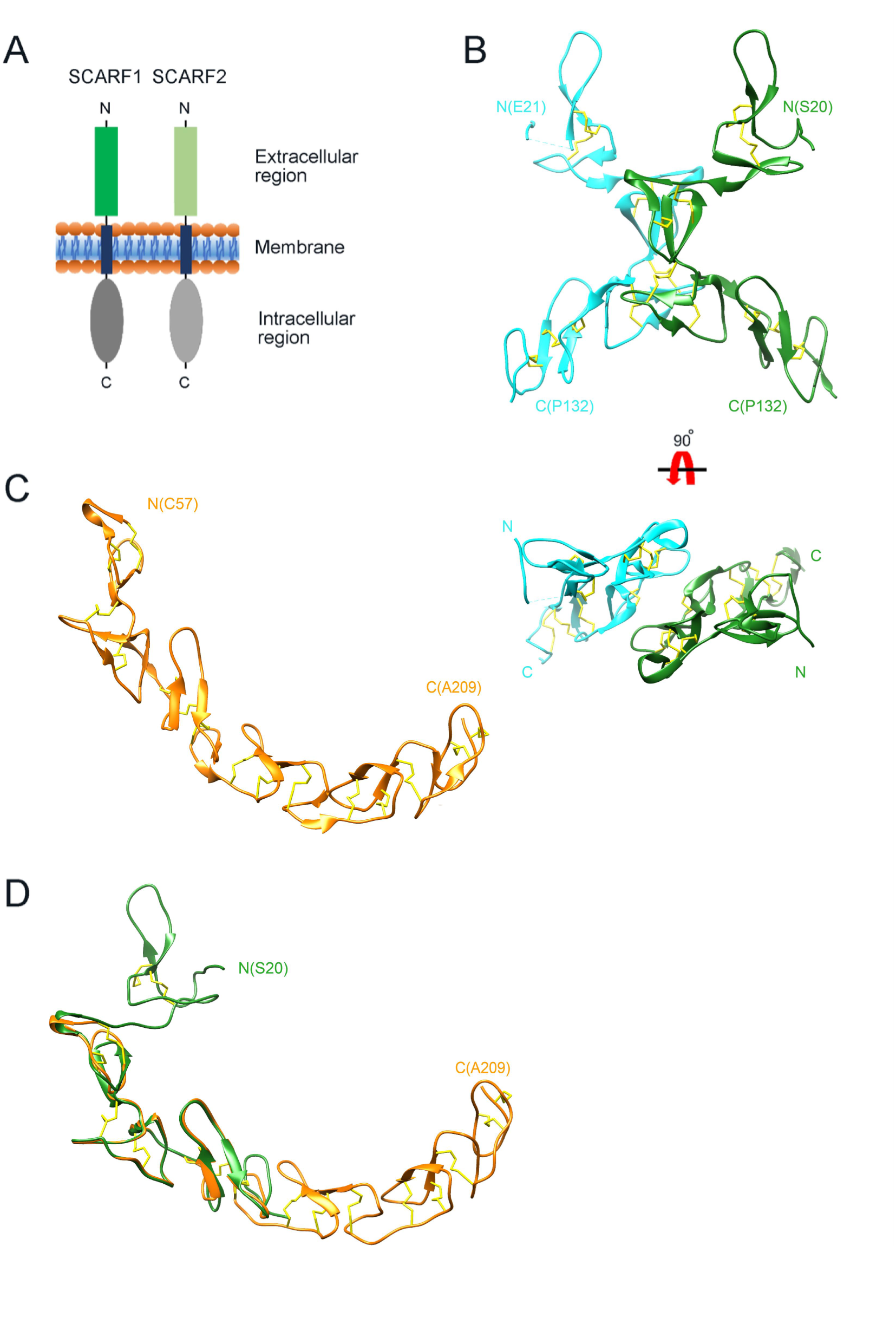
Crystal structure of the N-terminal fragments of SCARF1. (A) A schematic model of human SCARF1 and SCARF2. (B) Ribbon diagrams of a homodimer of an N-terminal fragment (f1, 20-132 aa; two monomers are shown in cyan and green, respectively) of SCARF1. (C) A ribbon diagram of an N-terminal fragment (f2, 57-209 aa, gold) of SCARF1. (D) Structure of the N-terminal fragment of SCARF1 (20-209 aa) by superimposing the crystal structures of f1 (green) and f2 (gold). Disulfide bonds are shown in yellow (B-D).

Previous reports have shown that SCARF1 can bind a number of endogenous ligands other than modified LDLs, including heat shock proteins [19, 20], calreticulin [29], Ecrg4 [30], Tamm-Horsfall protein [31] and apoptotic cells [23, 32], and mediate ligand internalization and transport [29, 33]. It can also bind bacterial, viral, and fungal antigens [7, 22, 34, 35], but the mechanisms for having such diverse ligand binding properties are unclear. By contrast, SCARF2 shows no binding activity with modified LDLs [24, 36], but recent data suggest SCARF2 may share ligands such as complement C1q and calreticulin with SCARF1 [36]. In addition, MEGF10 (multiple EGF-like domains-10), which might be another member of SR-F family, is a mammalian ortholog of Ced-1 and a receptor of amyloid-β in the brain [13, 37, 38], suggesting that SR-F family members may have rather wide ligand binding specificities.

SCARF1 (MW, 86 kD) is a type I transmembrane protein that has a short signal peptide followed by a long extracellular region, a transmembrane helix and a large cytoplasmic portion [14] (Fig. 1A). Its ectodomain has three glycosylation sites and contains multiple epidermal growth factor (EGF)-like domains, which usually has a two-stranded β-sheet followed by a loop region and three conserved disulfide bonds [14, 17, 39]. It has been shown that modified LDLs are the ligands for several SRs, including SR-A members, scavenger receptor class B type I (SR-BI), lectin-like oxidized low-density lipoprotein receptor (LOX-1), CD36, etc. [40–43]. For the SR-A members, including SCARA1 (CD204, SR-A1), MARCO and SCARA5, they bind modified LDLs in a Ca^2+^-dependent manner through the scavenger receptor cysteine-rich (SRCR) domains of the receptors [43]. In the case of LOX-1, both positively charged and non-charged hydrophilic residues might be involved in lipoprotein recognition [44, 45]. And for CD36, positively charged residues are identified to be critical for its binding with OxLDL [46]. Since SCARF1 has different structural features with other known lipoprotein receptors, how it recognizes modified lipoproteins remains unclear.

Here we determined the crystal structures of the N-terminal fragments of SCARF1 and characterized the interaction of SCARF1 with modified LDLs by biochemical and mutagenesis studies, thus providing mechanistic insights into the recognition of lipoproteins by this receptor.

## Results

### Crystal structures of the N-terminal fragments of SCARF1

To determine the structure of the extracellular region of SCARF1, the intact and several truncation fragments of SCARF1 ectodomain were expressed in insect cells and purified for crystallization screening. Among them, two fragments (f1, 20-132 aa and f2, 20-221 aa) (Fig. S5) were crystallized and the crystals diffracted to 2.2 Å and 2.6 Å, respectively (Table S1 and S2). The initial phasing for f1 crystal was done by SAD using Pt derivatives and the structure was refined to 2.2 Å with a native dataset after molecular replacement. The crystal of f1 fragment belongs to space group P212121 with two molecules per asymmetric unit (Fig. 1B). The f1 crystal structure contains a number of loops and two stranded β-sheets stabilized by hydrogen bonds and disulfide bonds, which is consistent with the typical feature of EGF-like domains, and it adopts a bow-like conformation with the middle part (∼55-102 aa) protruding outwards (Fig. 1B). The electron density of the N-terminal end (∼21-62 aa) of one molecule in an asymmetric unit is relatively weak and some residues are missing, probably due to the flexibility of this region.

The structure of f2 fragment was solved by molecular replacement using the structure of f1 fragment combined with the EGF-like fragments predicted by AlphaFold as phasing models and refined to 2.6 Å resolution. The crystal belongs to space group P4122 with one molecule per asymmetric unit (Fig. 1C). In the f2 crystal structure, the N-terminal region of SCARF1 (residue 20-56 aa) and the C-terminal region (210-221 aa) are largely missing, suggesting these regions are quite flexible, consistent with low electron density of the N-terminal end of the f1 crystal. Rest of the f2 fragment adopts a long-curved conformation with multiple EGF-like domains arranged in tandem (Fig. 1C). Superposition of the two crystal structures reveals the structure of the N-terminal fragment of SCARF1 (20-209 aa) (Fig. 1D), which has similar structural features with the AlphaFold prediction of the molecule (Fig. S1A). However, the crystal structures show a much larger curvature and local differences with the AlphaFold model (Fig. S1A, S2C), which is in agreement with a recent report regarding the comparison between the protein structures in PDB and AlphaFold models [47].

### SCARF1 forms homodimers

In the crystal of f1 fragment, two molecules in an asymmetric unit are related by a two-fold non-crystallographic symmetry axis. Interestingly, in the crystal of f2 fragment, although an asymmetric unit has only one molecule, the dimer related by the crystallographic two-fold symmetry can be well superimposed with the dimer found in the asymmetric unit of f1 crystals (Fig. 2A), suggesting that dimerization occurs in a similar fashion for the two fragments in different crystal forms. At the dimeric interfaces of f1 and f2 fragments, hydrogen bonds are formed between S88 and Y94 of the monomers, and F82 and Y94 are also close to each other, thus may form π-π interaction (Fig. 2B-C, S2A). Two salt bridges are also observed in f1 crystals between two monomers, one is between D61 and K71, the other is between R76 and D98 (Fig. 2B). But the salt bridges are not found in f2 crystals, suggesting they are not required for dimerization.

**Figure 2.**
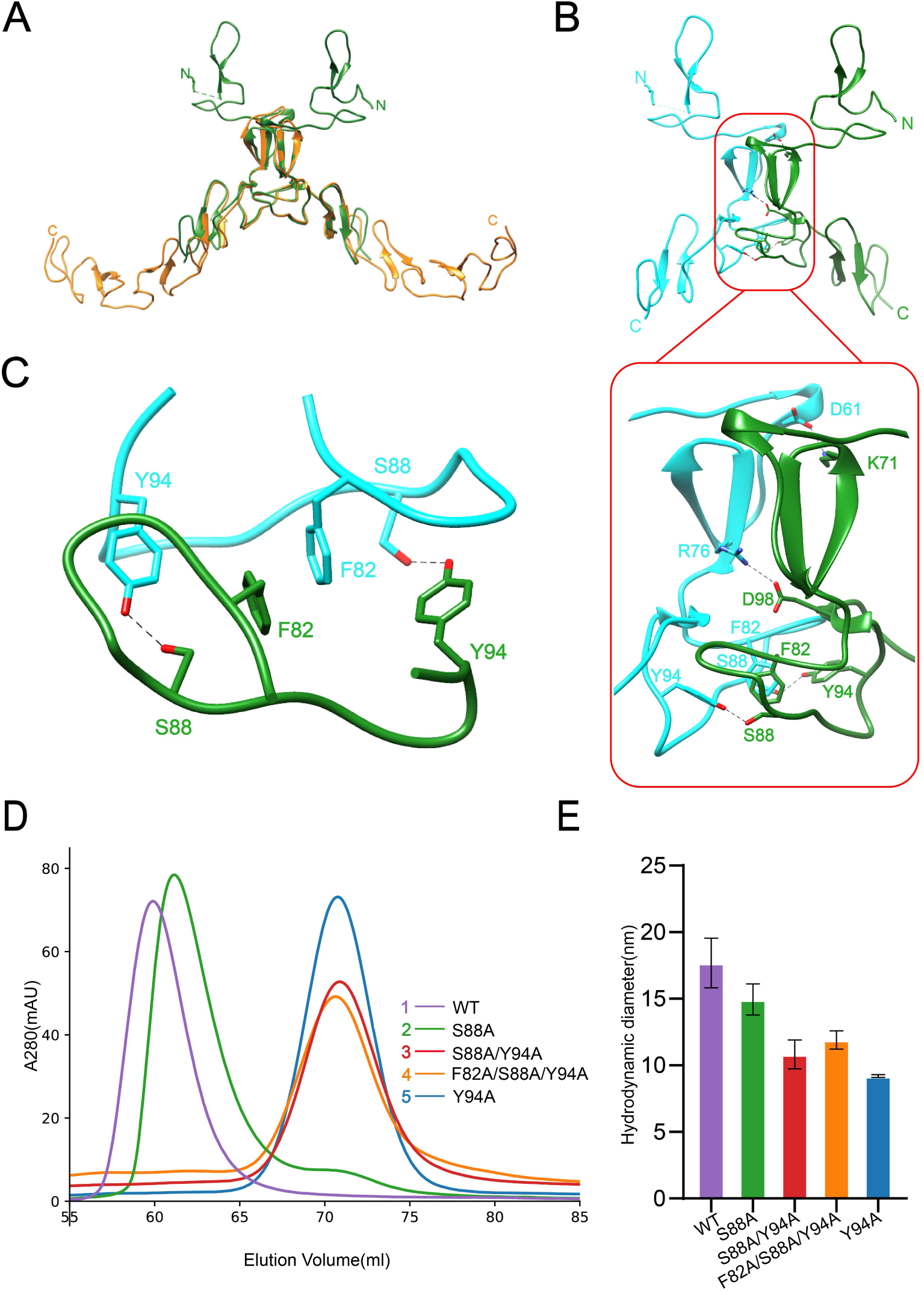
Dimerization of SCARF1. (A) Superposition of the homodimers of f1 (green) and f2 (gold) in the crystals. (B) The dimeric interface of f1 fragment of SCARF1 (red rectangles). Two monomers are colored in cyan and green, respectively. The side chains of the residues that form hydrogen bonds (dashed lines), salt bridges (dashed lines) and π-π interactions are labeled. (C) A local view of the dimeric interface of SCARF1. The side chains of the residues that form hydrogen bonds (dashed lines) and π-π interactions are labeled. (D) The SEC profiles of the wild type and the mutants of SCARF1 ectodomain. (E) The hydrodynamic diameters of the wild type and the mutants of SCARF1 ectodomain measured by DLS.

To further characterize the dimerization of SCARF1, a number of mutants of the ectodomain were constructed, and the size exclusion chromatography (SEC) was applied to monitor the elution volumes of the proteins (Fig. 2D, S3A). The results showed that mutant S88A, where the hydrogen bonds between S88 and Y94 were removed (Fig. 2C), had a small elution volume shift compared to the wild type (Fig. 2D), suggesting that the homodimers were still maintained, but instability may increase for the dimer. By contrast, mutant Y94A, where the hydrogen bonds and the π-π interactions between Y94 and F82 were both removed, showed a significant elution volume shift (Fig. 2D) and the volume corresponded to the molecular weight of the monomeric SCARF1 ectodomain, suggesting that the π-π interactions between F82 and Y94 were important for dimerization. Other mutants such as S88A/Y94A and F82A/S88A/Y94A also eluted at the volume of the monomer (Fig. 2D). In parallel, we measured the hydrodynamic diameters of the mutants by dynamic light scattering (DLS), and it showed that the wild type and mutant S88A had larger diameters than mutants Y94A, S88A/Y94A and F82A/S88A/Y94A (Fig. 2E, S3B), which was consistent with the SEC data. In addition, both the dimeric wild type and the monomeric mutant (S88A/Y94A) of the ectodomain exhibited similar SEC elution volumes at pH 6.0 and pH 8.0, suggesting that pH did not have large impact on the dimerization or the conformation of SCARF1 (Fig. S3C-D).

### Interactions of SCARF1 with modified LDLs

To characterize the interaction of SCARF1 with lipoproteins, we monitored the binding of lipoproteins with the SCARF1-transfected HEK293 cells using flow cytometry. The results showed that the SCARF1-transfected cells only bound OxLDL or AcLDL, rather than native LDL (Fig. 3A, S6), consistent with the previous reports [14, 48], and SCARF1 appeared to have higher binding activity for OxLDL than AcLDL (Fig. 3A). In parallel, we also tested the interaction of SCARF1 with high density lipoproteins (HDL) and oxidized HDL (OxHDL), the results showed that both HDL and OxHDL were not able to bind the SCARF1-transfected cells (Fig. 3A). Fluorescent confocal images also confirmed that SCARF1 colocalized with OxLDL or AcLDL, rather than LDL, HDL or OxHDL in the transfected cells (Fig. 3D). Moreover, the flow cytometry data showed that the binding between SCARF1 and modified LDLs occurred in the presence of Ca^2+^ or EDTA/EGTA (Fig. 3B-C, S7), suggesting that the interaction of SCARF1 with OxLDL or AcLDL is Ca^2+^-independent.

**Figure 3.**
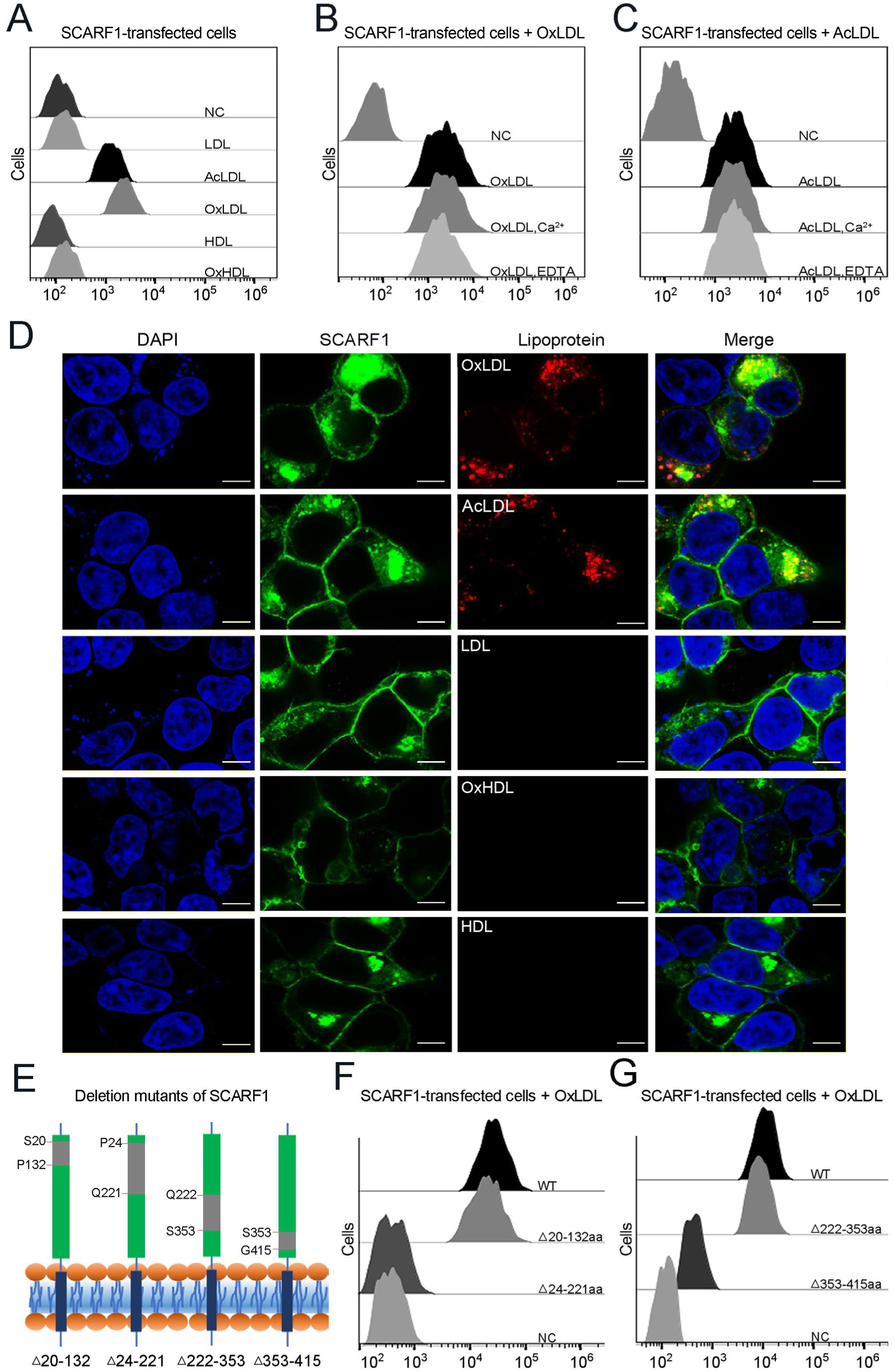
SCARF1 recognizes the modified LDLs. (A) Interactions of LDL, AcLDL, OxLDL, HDL and OxHDL with the SCARF1-transfected cells by flow cytometry (NC represents the non-transfected cells and no lipoprotein was added here). (B) Interactions of OxLDL with the SCARF1-transfected cells in the presence of Ca^2+^ or EDTA by flow cytometry (OxLDL was added for NC). (C) Interactions of AcLDL with the SCARF1-transfected cells in the presence of Ca^2+^ or EDTA by flow cytometry (AcLDL was added for NC). (D) Confocal fluorescent images of the SCARF1-transfected cells incubated with OxLDL, AcLDL, LDL, OxHDL or HDL (scale bar, 7.5 µm). (E) Schematic diagrams of the deletion mutants of SCARF1. The deleted regions are labeled and shown in gray. (F) Interactions of OxLDL with the deletion mutants SCARF1^Δ20-132aa^ and SCARF1^Δ24-221aa^ by flow cytometry (G) Interactions of OxLDL with the deletion mutants SCARF1^Δ222-353aa^ and SCARF1^Δ353-415aa^ by flow cytometry

To identify the lipoprotein binding region on the ectodomain of SCARF1, we generated a series of truncation mutants, including SCARF1^Δ20-132aa^, SCARF1^Δ24-221aa^, SCARF1^Δ222-353aa^ and SCARF1^Δ353-415aa^ (Fig. 3E). The cells transfected with these mutants were applied to examine the interactions with modified LDLs by flow cytometry. The results showed that the cells transfected with SCARF1^Δ20-132aa^ and SCARF1^Δ222-353aa^ could bind to OxLDL, similar to the cells expressing the wild type (Fig. 3F-G). By contrast, the cells expressing SCARF1^Δ24-221aa^ almost lost binding with OxLDL completely, suggesting that the binding site might locate at the middle region (133-221aa) of the ectodomain of SCARF1 (Fig. 3F). In addition, the cells expressing SCARF1^Δ353-415aa^ showed reduced binding with OxLDL (Fig. 3G), this may not be surprising as the deletion of the C-terminal regions of the ectodomain might change the conformation of the molecule and generate hinderance for the accessibility of lipoprotein particles.

### SCARF1 recognizes modified LDLs through charge interactions

To identify the binding site for modified LDLs on the region identified above (133-221aa), we calculated the surface electrostatic potential of the region based on the crystal structure of f2 fragment [49] and found a positively charged area in this region, which is mainly composed of two sites, one is R160 and R161 (Site 1), the other is R188 and R189 (Site 2) (Fig. 4A). To test whether these positively charged sites are involved in lipoprotein recognition, we generated a number of mutants of SCARF1 including: R160S, R161S and R160S/R161S for Site 1; R188S, R189S and R188S/R189S for Site 2; R160S/R161S/R188S, R160S/R188S/R189S, R160S/R161S/R188S/R189S for both sites, and monitored their binding with OxLDL or AcLDL using flow cytometry. The results showed that all the single mutants for the two sites R160S, R161S, R188S, R189S had similar binding activities as the wild type (Fig. 4B-C), whereas the two double mutants, R160S/R161S for Site 1 or R188S/R189S for Site 2, exhibited reduced binding with modified LDLs (Fig. 4D-E). And the double mutant for Site 2, R188S/R189S, appeared to have lower binding than the mutant for Site 1, R160S/R161S (Fig. 4D-E), implying that Site 2, R188S/R189S, may contribute more to the binding. The triple or quadruple mutants, R160S/R161S/R188S, R160S/R188S/R189S and R160S/R161S/R188S/R189S, where positive charges are largely removed for the two sites, almost lost binding activities with modified LDLs completely (Fig. 4D-E), suggesting these arginines are crucial for lipoprotein interaction and both sites contribute to the recognition. To further validate the importance of the charged residues, we made mutants where arginines were substituted with lysines on the two sites, including R160K/R161K, R188K/R189K and R160K/R161K/R188K/R189K. The flow cytometry data showed that all three mutants retained binding activities with modified LDLs (Fig. 4F-G), confirming the importance of charge interactions. Among them, mutants R160K/R161K and R188K/R189K had similar binding activities as the wild type, and mutant R160K/R161K/R188K/R189K exhibited a reduction in lipoprotein binding, suggesting the side chains of lysines may be slightly unfavorable for the interactions (Fig. 4F-G). In addition, fluorescent confocal images also confirmed that the mutant R160S/R161S/R188S/R189S did not bind to OxLDL, while the mutants R160K/R161K and R188K/R189K retained binding with OxLDL (Fig. 4H). Taken together, these data suggest that charge interactions are indispensable for the recognition of modified LDLs by SCARF1.

**Figure 4.**
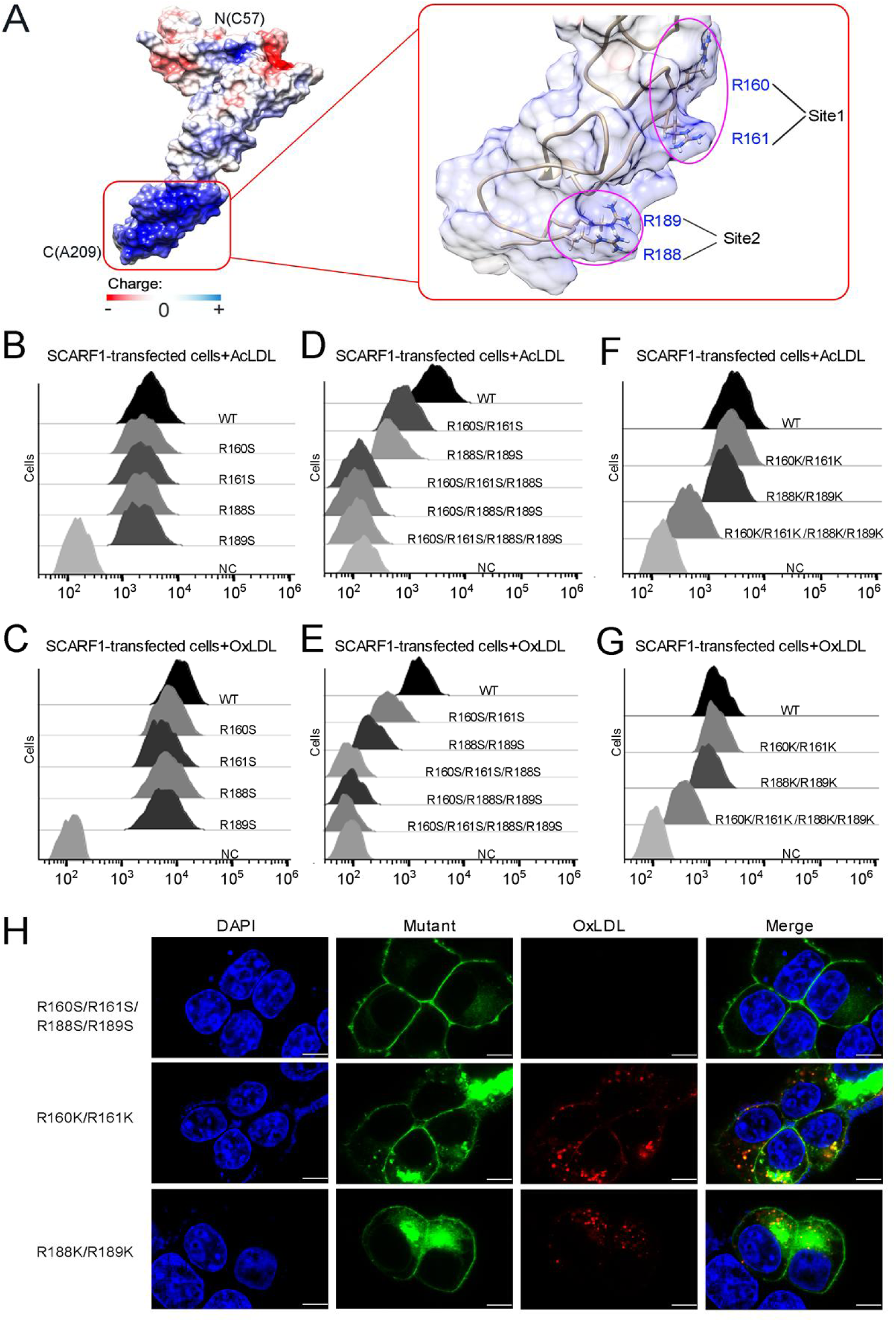
The binding sites of modified LDLs on SCARF1. (A) Surface electrostatic potential of f2 fragment shows a positively charged region on SCARF1 (left, red rectangle), which contains four arginines at site 1 and site 2 (right, magenta ovals). (B) Interactions of AcLDL with the cells transfected with the single mutants (R to S) of the binding sites by flow cytometry. (C) Interactions of OxLDL with the cells transfected with the single mutants (R to S) of the binding sites by flow cytometry. (D) Interactions of AcLDL with the cells transfected with the double, triple or quadruple mutants (R to S) of the binding sites by flow cytometry. (E) Interactions of OxLDL with the cells transfected with the double, triple or quadruple mutants (R to S) of the binding sites by flow cytometry. (F) Interactions of AcLDL with the cells transfected with the double or quadruple mutants (R to K) of the binding sites by flow cytometry. (G) Interactions of OxLDL with the cells transfected with the double or quadruple mutants (R to K) of the binding sites by flow cytometry. (H) Confocal fluorescent images of the SCARF1 mutant transfected cells incubated with OxLDL (scale bar, 7.5 µm).

To confirm the binding position identified above for modified LDLs, we expressed and purified the wild type and mutants of the ectodomain of SCARF1 for ELISA (Fig. 5A). The results were consistent with the flow cytometry data, showing that the mutant of the arginines, R160S/R161S/R188S/R189S, lost binding activity with OxLDL (Fig. 5A). The ELISA data also showed that the monomeric mutants (S88A/Y94A, F82A/S88A/Y94A) had slightly higher affinities with OxLDL than the dimeric wild type, which might be due to the steric hinderance of the dimers when OxLDLs were coated onto the plates (Fig. 5A), but flow cytometry suggested that the monomeric mutants had lower binding than the wide type (Fig. 5B), implying that the dimeric form may be more efficient to recognize lipoproteins on the cell surface. In addition, we also tested the binding of SCARF1 with OxLDL at pH 6.0 by ELISA and flow cytometry, and both data suggested that the binding activity was retained at pH 6.0 (Fig. 5A-B).

**Figure 5.**
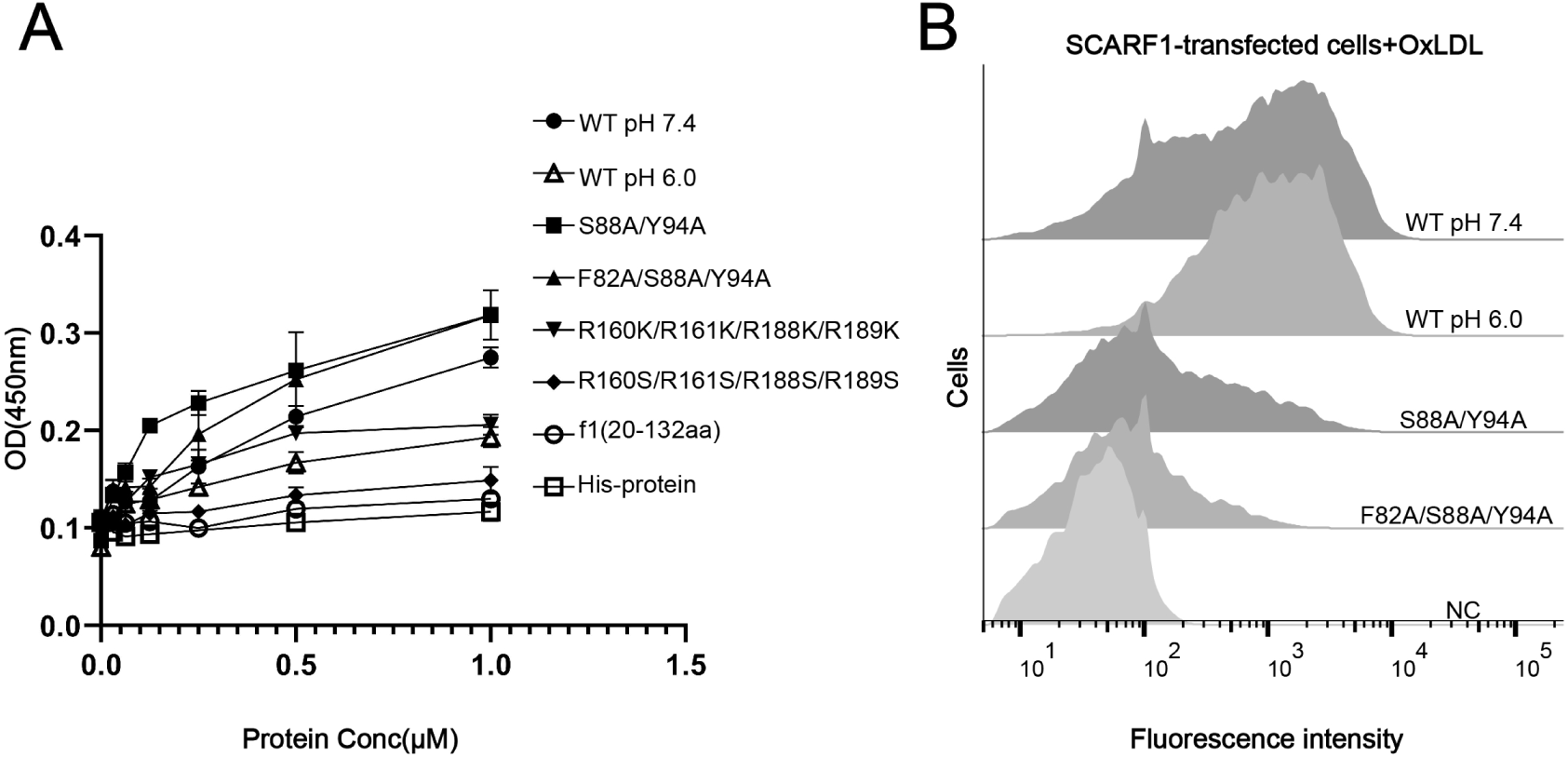
The binding of SCARF1 with OxLDL. (A) ELISA of the interactions of OxLDL with the wild type and the mutants of the ectodomain of SCARF1(f1 fragment is also applied). The assays are performed at pH 7.4 if not labeled. The ectodomain of human HER3 (His-protein) is applied as a control. (B) Interactions of OxLDL with the cells transfected with the wild type and the mutants of SCARF1 by flow cytometry. The assays are performed at pH 7.4 if not labeled.

### Interactions of modified LDLs with SCARF1/SCARF2 chimeric molecules

As another member in SR-F class, SCARF2 has similar structural features with SCARF1 according to AlphaFold prediction (Fig. S1A-B). However, the flow cytometry data showed that SCARF2 did not bind to AcLDL or OxLDL (Fig. 6A-B), which is in agreement with the previous reports [24, 36]. To further validate the lipoprotein binding sites identified on SCARF1, we generated three pairs of SCARF1/SCARF2 chimeric molecules by switching the counterparts of the two molecules, including: i) SF1-1 and SF2-1, where 1-421aa of SCARF1(ectodomain) and 1-441aa of SCARF2 (ectodomain) are switched; ii) SF1-2 and SF2-2, where 1-221aa of SCARF1 and 1-242aa of SCARF2 are switched; and iii) SF1-3 and SF2-3, where 133-221aa of SCARF1 and 156-242aa of SCARF2 are switched (Fig. 6C-E). Both flow cytometry data and fluorescent confocal images showed that SF2-1 gained binding activity with modified LDL, whereas SF1-1 lost the affinity, suggesting that the ectodomain of SCARF1 is sufficient for lipoprotein interaction (Fig. 6C, 6F-G). Furthermore, the flow cytometry results of SF1-2, SF2-2, SF1-3 and SF2-3 demonstrated that SCARF2 chimeric molecules could bind modified LDLs when its counterparts are replaced by the fragments from SCARF1 that contain the binding sites (Fig. 6D-F), thereby confirming the lipoprotein binding sites identified on SCARF1. However, the chimeric molecules SF2-2 and SF2-3 showed weaker affinities with OxLDL than SCARF1 or SF2-1 (Fig. 6F), suggesting the overall conformation or residues around the substituted binding region of SCARF2 might also affect ligand interaction. In addition, we made a sequence alignment of SCARF1 from different species and found that the positively charged residues at both site 1 and site 2 are well conserved, but not for SCARF2 (Fig. 6H, S2B-D), consistent with the importance of these charged residues in ligand recognition for SCARF1.

**Figure 6.**
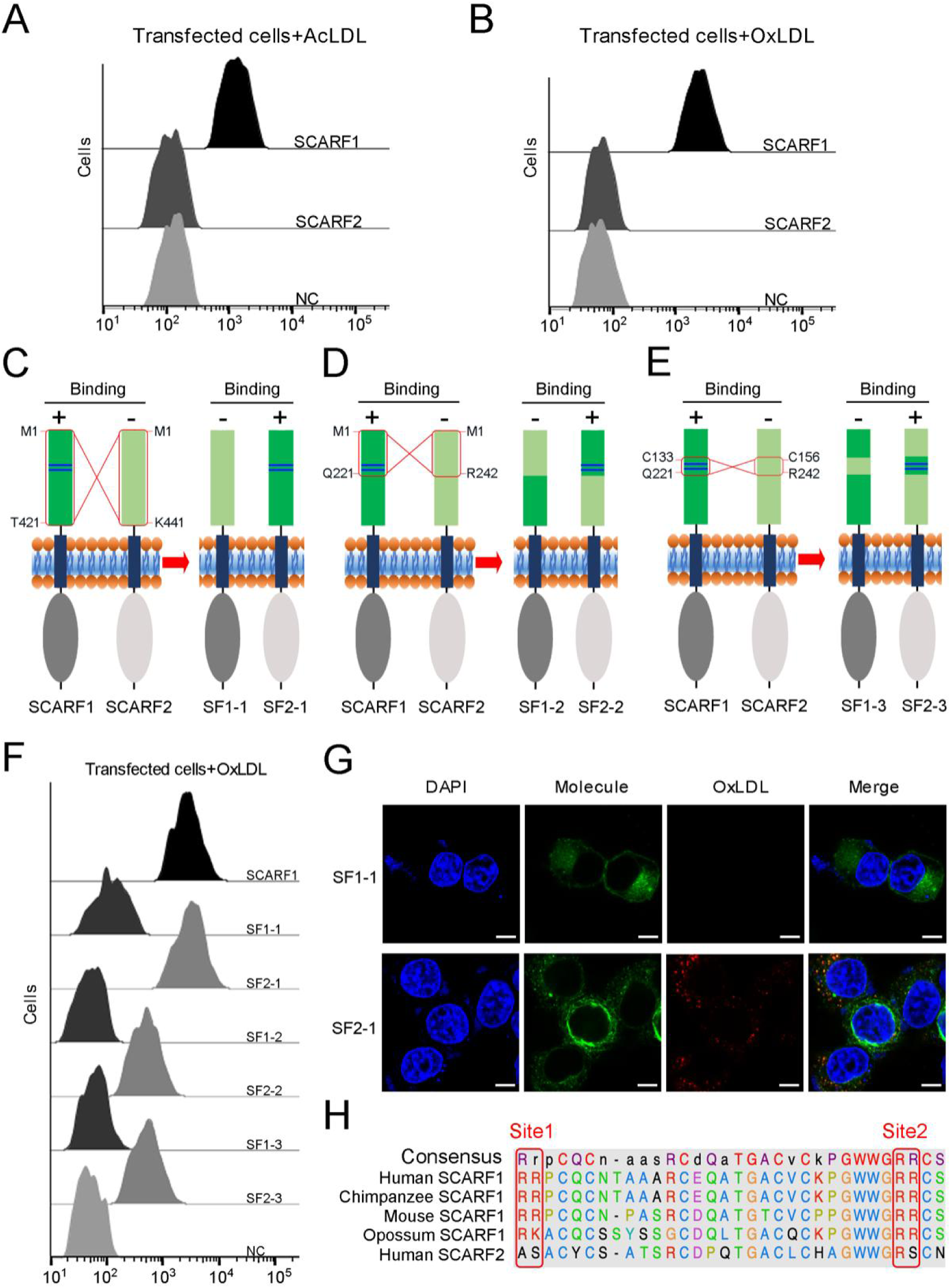
The interactions of modified LDLs with SCARF1-SCARF2 chimeric molecules. (A) Interaction of AcLDL with the SCARF1 or SCARF2 transfected cells by flow cytometry. (B) Interaction of OxLDL with the SCARF1 or SCARF2 transfected cells by flow cytometry. (C), (D) and (E) Schematic diagrams of the SCARF1-SCARF2 chimeric molecules generated for binding assays. The switched regions are indicated by red rectangles. The positively charged site 1 and site 2 of SCARF1 are shown as blue lines. The binding of molecules with OxLDL is indicated as + (positive) or – (negative). (F) Interactions of OxLDL with the chimeric molecule transfected cells by flow cytometry. (G) Confocal fluorescent images of the chimera molecule SF1-1 or SF2-1 transfected cells incubated with OxLDL (scale bar, 7.5 µm). (H) Sequence alignment of the binding sites of SCARF1 from different species and human SCARF2.

### Interaction of SCARF1 with modified LDLs is inhibited by teichoic acids

Previous reports have shown that SCARF1 has multiple ligands [14, 29-31, 50]. Among them, teichoic acids from *Staphylococcus aureus* has been shown to be able to bind SCARF1 in a charge-dependent manner and mediate adhesion to nasal epithelial cells *in vitro*, and the binding site of teichoic acids locates at the middle region (137-250 aa) of the SCARF1 ectodomain [51]. Here we found that the binding of SCARF1 with OxLDL or AcLDL could be inhibited by teichoic acids efficiently (Fig. 7A-B). And the two double mutants of SCARF1, where the lipoprotein binding sites are mutated, showed more inhibitory effects from teichoic acids (Fig. 7A-B), suggesting that the binding sites for modified LDLs and teichoic acids might be shared or overlapped with each other on the ectodomain of SCARF1.

**Figure 7.**
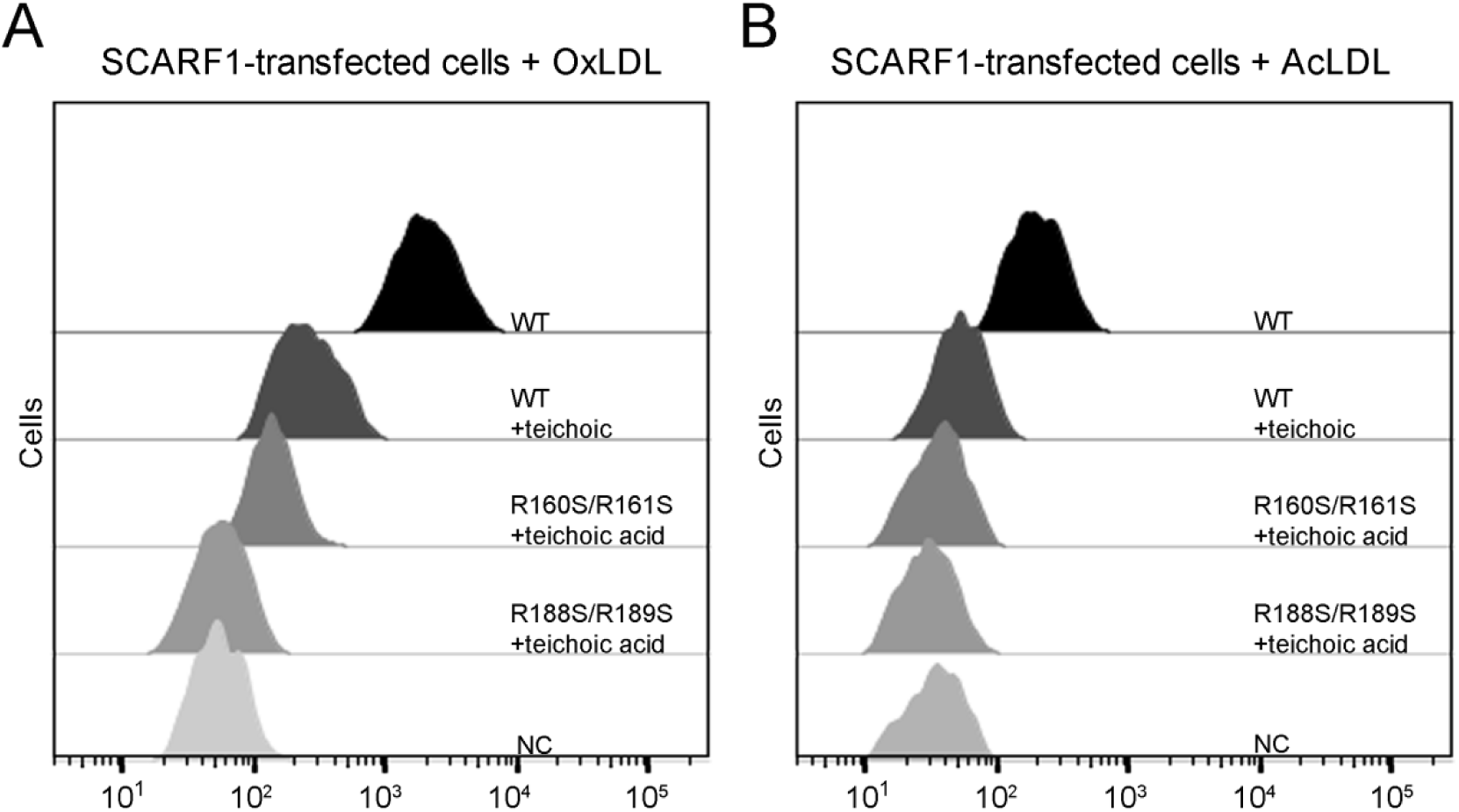
Inhibition of the interactions of SCARF1 with modified LDLs by teichoic acids. (A) Interaction of OxLDL with the cells transfected with the wild type or mutants of SCARF1 in the presence of teichoic acids by flow cytometry. (B) Interaction of AcLDL with the cells transfected with the wild type or mutants of SCARF1 in the presence of teichoic acids by flow cytometry.

## Discussion

SCARF1 can bind both endogenous and exogenous ligands through its ectodomain [52, 53], which adopts a long-curved conformation with multiple EGF-like domains arranged in tandem according to the crystal structures and the AlphaFold model. EGF-like domains are commonly found in cell surface molecules [54, 55] and usually bind ligands in a Ca^2+^-dependent manner [56–58]. The EGF-like domains of SCARF1 do not contain the typical Ca^2+^-binding sites according to sequence analysis [48] and no obvious Ca^2+^ density is observed in the crystal structures, consistent with the results of biochemical assays, suggesting that the interaction of SCARF1 with modified LDLs is Ca^2+^-independent. This is in contrast to the SR-A family, where SCARA1, MARCO and SCARA5 bind modified LDLs in a Ca^2+^-dependent manner [43].

The lipoprotein binding sites of SCARF1 locate at the middle region of the ectodomain and arginines are crucial for recognizing modified LDL by providing positive charges, which may have similarities with WIF-1, a Wnt inhibitory factor containing EGF-like domains that bind glycosaminoglycans through conserved arginines and lysines [59]. In fact, the binding of lipoproteins via charge interactions has been reported before. For LOX-1, which is also a scavenger receptor, positively charged residues such as arginines play essential roles for its binding with OxLDL [44, 45, 60]. And similarly, positively charged residues are important to recognize OxLDL by CD36, which is a member in scavenger receptor class B [46, 61]. Therefore, the interaction of SCARF1 with modified LDLs might be similar to LOX-1 or the SR-B members, rather than the SR-A members. Moreover, both structural and biochemical data suggest that SCARF1 forms homodimers (Fig. 8), which may facilitate the accessibility or binding of modified lipoproteins on the cell surface, and is also analogous to some of the lipoprotein receptors such as LOX-1[62, 63].

**Figure 8.**
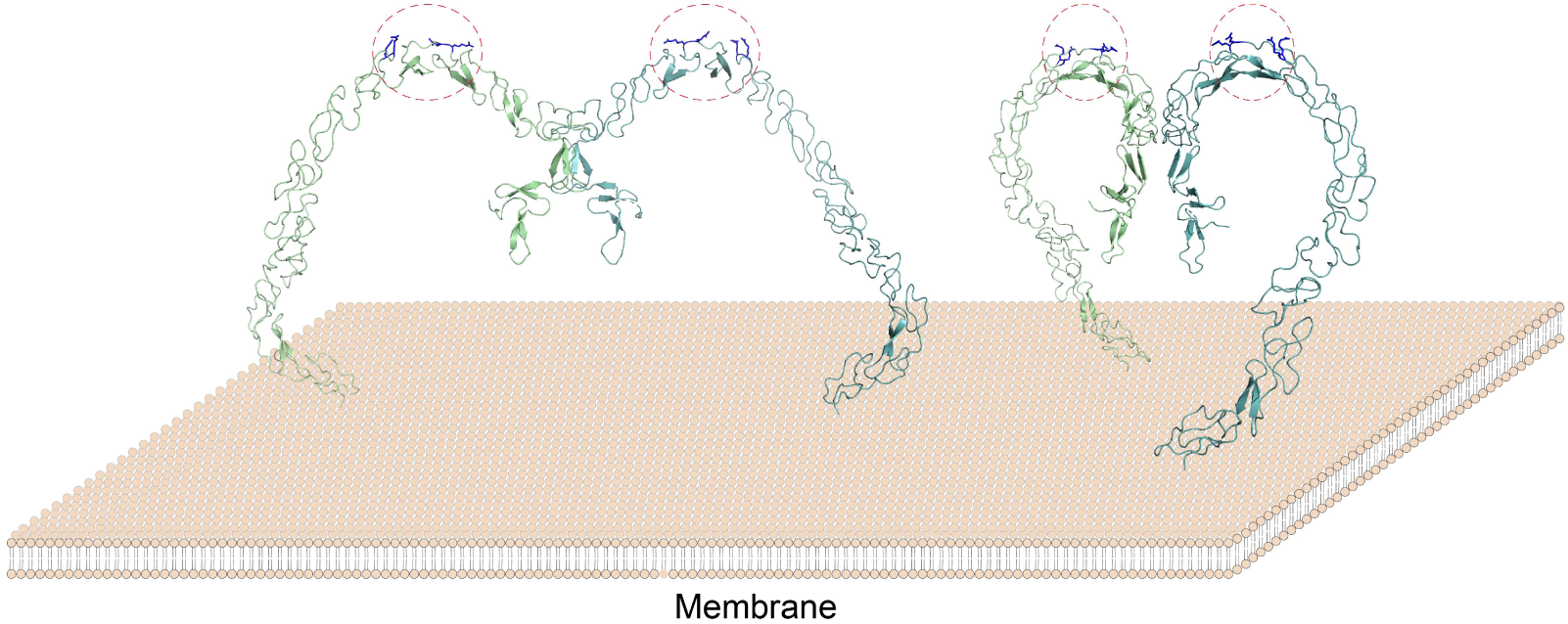
A model of the SCARF1 homodimers on the membrane surface. The structure of the SCARF1 homodimer is generated by combining the crystal structures of f1 and f2, and rest of the ectodomain is from AlphaFold prediction. Two dimers are shown with different viewing angles. The monomers are colored in green or cyan. The lipoprotein binding sites are labeled by red dashed ovals and the arginines at the binding sites are colored in blue.

Among the ligands of SCARF1, the teichoic acids from *Staphylococcus aureus* have been shown to interact with SCARF1 in a charge-dependent manner [51], which is similar to modified LDLs that also need charged residues to bind SCARF1. Therefore, teichoic acids and modified LDLs may share the binding sites on the ectodomain of SCARF1, implying that this region might be a general binding position for some of the scavenging targets, but whether all the ligands utilize this region for binding needs further investigation. In addition, the sequence alignment also shows that these charged residues are well conserved in different species for SCARF1, but SCARF2 lacks such sites on the ectodomain, suggesting SCARF1 and SCARF2 are functionally divergent, although they have some sequence similarities.

Over the past decades, SCARF1 has been linked to a number of diseases, including cardiovascular diseases, systemic lupus erythematosus, fungal keratitis and cancer [26, 32, 53, 64]. But the mechanisms and roles of SCARF1 in these diseases are largely elusive. The structural and mechanistic characterization of SCARF1 and its interactions of multiple ligands would shed light on the physiological and pathological roles of this receptor on the corresponding pathways and may also provide insights on the therapeutic strategies against the related diseases.

## Materials and methods

### Protein expression and purification

For protein expression in insect cells, the cDNA of human SCARF1 encoding f1 (20-132aa) and f2 (20-221aa) fragments were sub-cloned into a pFastBac vector fused with an N-terminal melittin signal peptide and a C-terminal 6xHis tag using NovoRec recombinant enzyme. Sf9 cells were used for generating recombinant baculoviruses and high five cells were used for protein production. The infected cells were cultured in ESF921 medium (Expression Systems) for 3 days in a 27°C humidified incubator. The supernatants were buffer-exchanged with 25 mM Tris, 150 mM NaCl at pH 8.0 by dialysis, then applied to Ni-NTA chromatography (Ni-NTA Superflow, Qiagen). The eluted proteins were concentrated to 1 ml using Amicon Ultra-15 3k-cutoff filter (Millipore) and loaded onto a HiLoad Superdex 75 prep grade column or a HiLoad Superdex 200 prep grade column (GE Healthcare) with Tris-NaCl buffer (10 mM Tris, 150 mM NaCl at pH 7.5) for further purification. The purified proteins were loaded onto SDS-PAGE (12%) and stained with coomassie brilliant blue R250 for detection.

For protein expression in mammalian cells, the cDNAs of the human SCARF1 ectodomain and mutants (S88A, Y94A, S88A/Y94A, F82A/S88A/Y94A) were sub-cloned into a pTT5 vector fused with a C-terminal 6xHis tag. The transfected HEK293F cells were cultured in suspension for 5 days in a CO_2_ humidified incubator at 37°C, then the supernatants were collected and applied to Ni Smart Beads (Smart-lifesciences). The eluted proteins were concentrated to 1 ml using Amicon Ultra-15 10k-cutoff filter (Millipore) and loaded onto a HiLoad Superdex 200 prep grade column (GE Healthcare) with Tris-NaCl buffer (50 mM Tris, 150 mM NaCl at pH 8.0) for further purification. The purified proteins were loaded onto SDS-PAGE (10%) and stained with coomassie brilliant blue R250 for detection.

### Cell culture and transfection

HEK293T cells were cultured in Dulbecco’s modified Eagle’s medium (HyClone) supplemented with 10% fetal bovine serum and cells were incubated in a humidified incubator at 37 °C with 5% CO_2_. The cells were seeded in a 12-well plate in advance, and on the day of transfection, the cells that were 80%-90% confluent in the 12-well plate were transfected using PEI (Polyplus). After 4-6 h of transfection, the culture medium was replaced by the fresh medium. Then about 24 h after transfection, the cells were ready for assays.

HEK293F cells were cultured in suspension with Union-293 medium (Union Bio) in a humidified incubator at 37 °C with 8% CO_2_. Transfections were done at the cell density of 1.5-2.0×10^6^/ml using PEI (Polyplus) and after 24 h, the cells were ready for assays.

### Crystallization and structural determination

The purified protein was concentrated to 30 mg/ml (f1, 20-132aa) and 10 mg/ml (f2, 20-221aa) (measured by UV absorption at 280nm) using Amicon Ultra-15 3k-cutoff filter (Millipore). Crystal screening was done at 18 °C by sitting-drop vapor diffusion method using 96-well plates (Swissci) with commercial screening kits (Hampton Research). A Mosquito nanoliter robot (TTP Labtech) was used to set up 200 nl protein sample mixed with 200 nl reservoir solution. The crystals of f1 were grown in a solution containing 20% (w/v) polyethylene glycol 3350, 0.2 M ammonium sulfate, 0.1 M Bis-Tris (pH 5.5) and the crystals of f2 were grown in 0.1 M citric acid/sodium citrate (pH 3.6), 0.2 M sodium citrate tribasic, 9% PEG6000. Crystals of both fragments were obtained after 5-7 days. 10% ethylene glycol was added during crystal harvest and data collection as cryo-protectant. The f1 crystal heavy atom derivatives were obtained by soaking with 10 mM K_2_PtCl_4_ for 2 minutes before data collection. Diffraction data were collected using a PILATUS 6M detector at BL18U1 beamline of National Facility for Protein Science Shanghai (NFPS) at Shanghai Synchrotron Radiation Facility (SSRF). X-ray diffraction data were integrated and scaled with HKL-3000 package [65].

The initial phasing of the f1 crystals was done by SAD with Pt derivatives, then the structure was used as a search model of molecular replacement for a native data set of f1 with higher resolution for further refinement. The f2 crystal structure was solved by molecular replacement using the structure of f1 fragment combined with the EGF-like fragment predicted by AlphaFold as search models using Phenix [66, 67]. Model building and refinement were done using Coot [68] and Phenix [69, 70]. The structural figures were prepared using UCSF Chimera [71]. The coordinates and the structure factors have been deposited in the Protein Data Bank with entry 8HN0 and 8HNA for f1 and f2 fragments, respectively.

### Mutagenesis experiments

SCARF1 mutations, including base-substitution mutations (R160S, R161S, R188S, R189S, R160S/R161S, R188S/R189S, R160K/R161K, R188K/R189K, R160S/R161S/R188S, R160S/R188S/R189S, R160S/R161S/R188S/R189S, R160K/R161K/R188K/R189K,S88A,Y94A, S88A/Y94A, F82A/S88A/Y94A), deletion mutations (SCARF1^Δ20-132aa^, SCARF1^Δ24-221aa^, SCARF1^Δ222-353aa^, SCARF1^Δ353-415aa^) were introduced into the corresponding plasmids by PCR using KOD DNA polymerase (Sparkjade Science Co., Ltd.). The template plasmids were digested by DpnI (Thermo Fisher Scientific), and the digested PCR products were ligated by Ligation High (TOYOBO). The chimeric molecules of SCARF1 and SCARF2 were constructed by replacing fragments (1-441aa, 1-242aa, or 156-242aa) of SCARF2 with its counterparts of SCARF1 (1-421aa, 1-221aa or 133-221aa), respectively.

### Dynamic Light Scattering

The purified protein samples were concentrated to about 100 µg/mL in Tris buffer (50 mM Tris, 150 mM NaCl at pH 8.0) and dynamic light scattering signals were measured and processed on a Zetasizer Pro analyzer (Malvern Panalytical). Data for each sample were collected in triplicate at 25°C.

### Preparation of modified lipoproteins

Lipoproteins (purity, 97%-98%), including Dil-LDL (20614ES76), Dil-AcLDL (20606ES76), Dil-OxLDL (20609ES76), LDL (20613ES05), AcLDL (20604ES05) were purchased from Yeasen. Dil-HDL (H8910) was purchased from Solarbio. The lipoproteins mentioned above (LDL, AcLDL, OxLDL, HDL) were isolated from human plasma.

For preparation of OxHDL, 140 µl of 1 mg/ml HDL were buffer-exchanged to PBS solution, and then the same volume of 100 µM CuSO_4_ was divided into multiple small portions and added to HDL solutions gradually. After 16 h at 37°C, the reaction solutions were dialyzed against the PBS buffer containing 0.3 mM EDTA to stop the reaction.

### Flow cytometry

For the binding assays of SCARF1 and SCARF2 with lipoproteins, HEK293T cells were transiently transfected with the full-length SCARF1 or SCARF2 fused with a C-terminal GFP tag. After 24 h, 5 µg Dil-tagged (wavelength: 565 nm) lipoprotein (Dil-LDL, Dil-AcLDL, Dil-OxLDL, Dil-HDL, Dil-OxHDL) was added to the culture medium. After 2 to 4 h at 37°C, cells were washed three times with the washing buffer (25 mM Hepes, 150 mM NaCl, 0.1% Tween 20, pH 7.4) and then washed twice with the cleaning buffer (25 mM Hepes, 150 mM NaCl, pH 7.4) for flow cytometry.

For the Ca^2+^ assays, HEK293T cells were transiently transfected with the full-length SCARF1 fused with a C-terminal GFP tag. After 24 h, 5 µg Dil-labeled lipoprotein (Dil-AcLDL, Dil-OxLDL) was added to the culture medium containing 2 mM Ca^2+^ or 2 mM EDTA/EGTA. After 2 to 4 h, the cells were washed twice with the corresponding washing buffer (25 mM Hepes, 150 mM NaCl, 0.1% Tween 20, 2 mM Ca^2+^, or 2 mM EDTA/EGTA, pH 7.4) and then washed twice with the cleaning buffer (25 mM Hepes, 150 mM NaCl, 2 mM Ca^2+^, or 2 mM EDTA/EGTA, pH 7.4) for flow cytometry.

For the binding assays of the SCARF1 mutants with lipoproteins, HEK293T cells were transiently transfected with the wild type or the mutants of SCARF1 fused with a C-terminal GFP tag. After 24 h, 5 µg Dil-tagged (wavelength: 565 nm) lipoprotein (Dil-LDL, Dil-AcLDL, Dil-OxLDL) was added to the culture medium. After 2 to 4 h at 37°C, cells were washed three times with the washing buffer (25 mM Hepes, 150 mM NaCl, 0.1% Tween 20, pH 7.4) and then washed twice with the cleaning buffer (25 mM Hepes, 150 mM NaCl, pH 7.4) for flow cytometry. For the assays performed at pH 6.0, PBS buffer (pH 6.0) was used following the similar procedure.

For the teichoic acids assays, HEK293T cells were transiently transfected with the full-length SCARF1 fused with a C-terminal GFP tag. After 24 h, teichoic acids (100 µg/ml) were added to the culture medium. After 1 to 2 h at 4°C, 5 µg Dil-labeled lipoprotein (Dil-AcLDL, Dil-OxLDL) was added to the culture medium. After 2 to 4 h at 37°C, cells were washed three times with the washing buffer (25 mM Hepes, 150 mM NaCl, 0.1% Tween 20, pH 7.4) and then washed twice with the cleaning buffer (25 mM Hepes, 150 mM NaCl, pH 7.4) for flow cytometry.

For the assays to monitor the expression of SCARF1 and mutants, HEK293F cells were transiently transfected with the wild type or the mutants fused with a C-terminal GFP tag. After 24 h, Human SREC-I/SCARF1 Alexa Fluor 647-conjugated Antibody ( FAB2409R, R&B SYSTEMS) was added to the culture medium. After incubation of 30 min at 4°C, cells were washed three times with the washing buffer (135 mM NaCl, 2.7 mM KCl, 1.5 mM KH_2_PO4, and 8 mM K_2_HPO4) for flow cytometry (Fig. S4).

Flow cytometry data were acquired using a LSR Fortessa flow cytometer (BD Biosciences). Data analysis was performed using FlowJo software (Tree Star, Inc).

### ELISA experiments

Lipoprotein (OxLDL) was coated onto 96-well plates with 1 μg protein per well at 4 °C overnight. The plates were then blocked with the blocking buffer (25mM Hepes,150mM Nacl,0.1% Triton X-100,and 5%(w/v) BSA, pH 7.4) at room temperature for 3 h. The purified His-tagged proteins was serially diluted and added to each well in the binding buffer (25mM Hepes,150mM Nacl,0.1% Triton X-100,and 2 mg/ml BSA, pH 7.4). After incubation at room temperature for 2 h, the plates were washed five times with the washing buffer (25mM Hepes,150mM NaCl and 0.1% Triton X-100, pH 7.4) and then incubated with the mouse anti-His antibody (66005-1-Ig, Proteintech) for 1 h. After washing three times with the washing buffer, the plates were incubated with the HRP-labeled goat anti-mouse IgG (A0216, Beyotime) for 1 h. After washing three times with the washing buffer, 100 μl TMB Chromogen Solution (P0209-500ml, Beyotime) was added to each well and incubated for 30 min at room temperature. Then, 50 μl H_2_SO_4_ (2.0M) was added to each well to stop the reactions. For the binding of lipoproteins at pH 6.0, PBS buffer (pH 6.0) was used following the similar procedure. The plates were then read at 450nm on a Synergy2 machine (Bio Tek Instruments).

### Confocal microscopy

HEK293T cells were grown on coverslips and transfected with the full-length SCARF1, SCARF2 or the mutants of SCARF1 fused with a C-terminal GFP tag using 6-well plates. After 24 h of transfection, 5 μg Dil-labeled Lipoprotein (Dil-LDL, Dil-AcLDL, Dil-OxLDL, Dil-OxHDL) added to the plates. After 2 to 4 h of incubation, the cells were fixed by 4% paraformaldehyde in TBS (50 mM Tris and 150 mM NaCl, pH7.4). After washing twice with the buffer (25 mM Hepes, 150 mM NaCl, 0.05% Tween-20, pH 7.4). Then the cells were blocked in the blocking buffer (25 mM Hepes, 150 mM NaCl, 5% (w/v) BSA, 0.05% Tween-20, pH7.4) at room temperature for 1 h. After washing three times with the washing buffer (25 mM Hepes, 150 mM NaCl, 0.05% Tween-20, pH 7.4) and the cells were incubated with 5 μM DAPI for 15 min. Then the plates were washed again for confocal microscopy with a Leica SP8 microscope.

## Data availability

The crystal structures of the human SCARF1 fragments, f1 and f2, are deposited in PDB (www.rcsb.org) with entry 8HN0 and 8HNA, respectively.

## Acknowledgements

We thank the Integrated Laser Microscopy System and the Large-scale Protein Production System at the National Facility for Protein Science in Shanghai (NFPS), Shanghai Advanced Research Institute, Chinese Academy of Sciences, China, for technical support. We also thank the beamline BL18U1 of National Facility for Protein Science Shanghai (NFPS) at Shanghai Synchrotron Radiation Facility for their assistance in X-ray diffraction data collection. This work is supported by National Natural Science Foundation of China (No. 91957102) to Y.H. and we also thank the support from Innovative research team of high-level local universities in Shanghai (SHSMU-ZLCX20212601).

## Supplementary Figures

**Figure S1.**
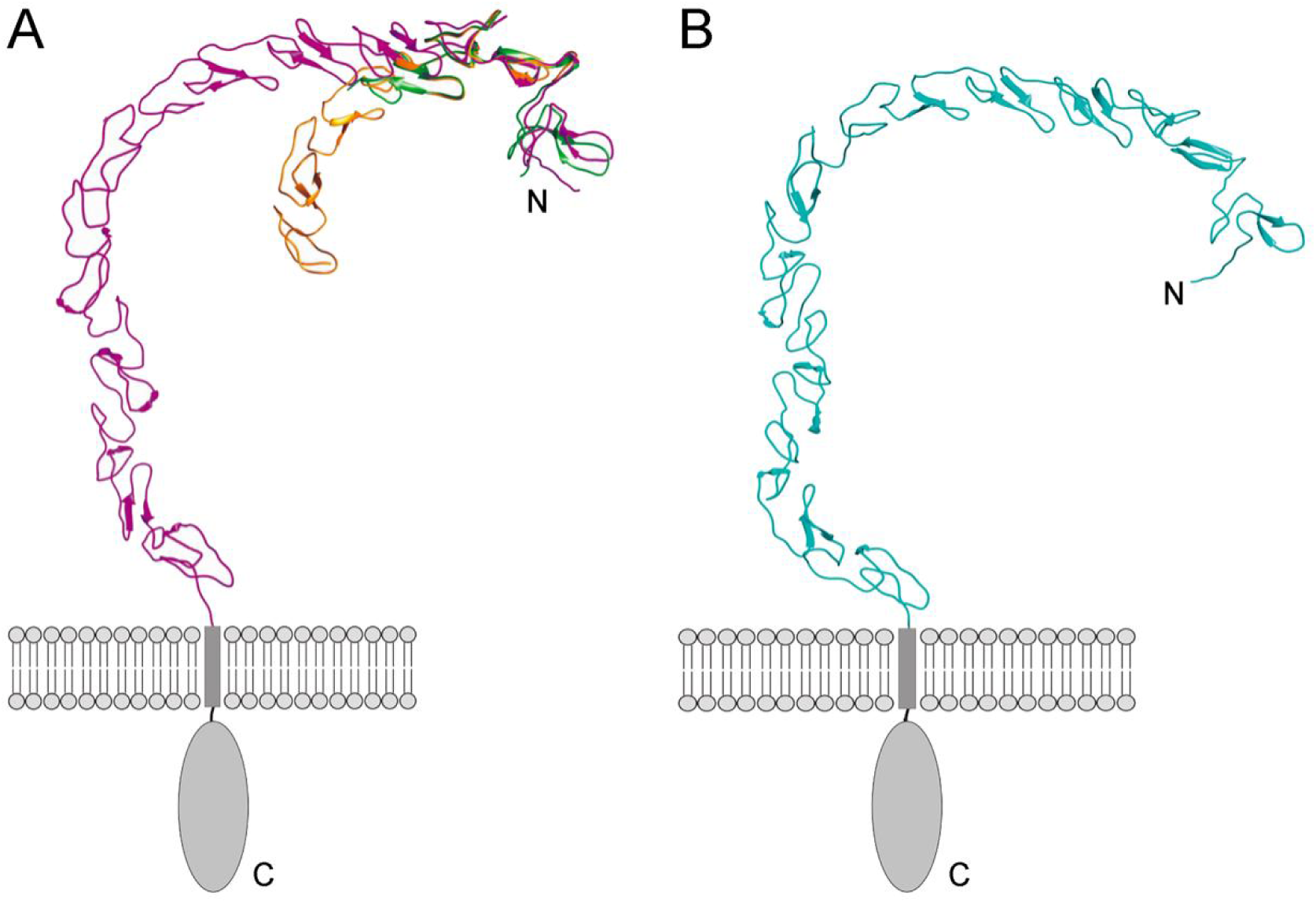
Models of the ectodomains of SCARF1 and SCARF2 by AlphaFold. (A) Superposition of the crystal structures of f1 (green) and f2 (gold) with the AlphaFold model of the SCARF1 ectodomain (magenta). (B) AlphaFold model of the SCARF2 ectodomain (cyan).

**Figure S2.**
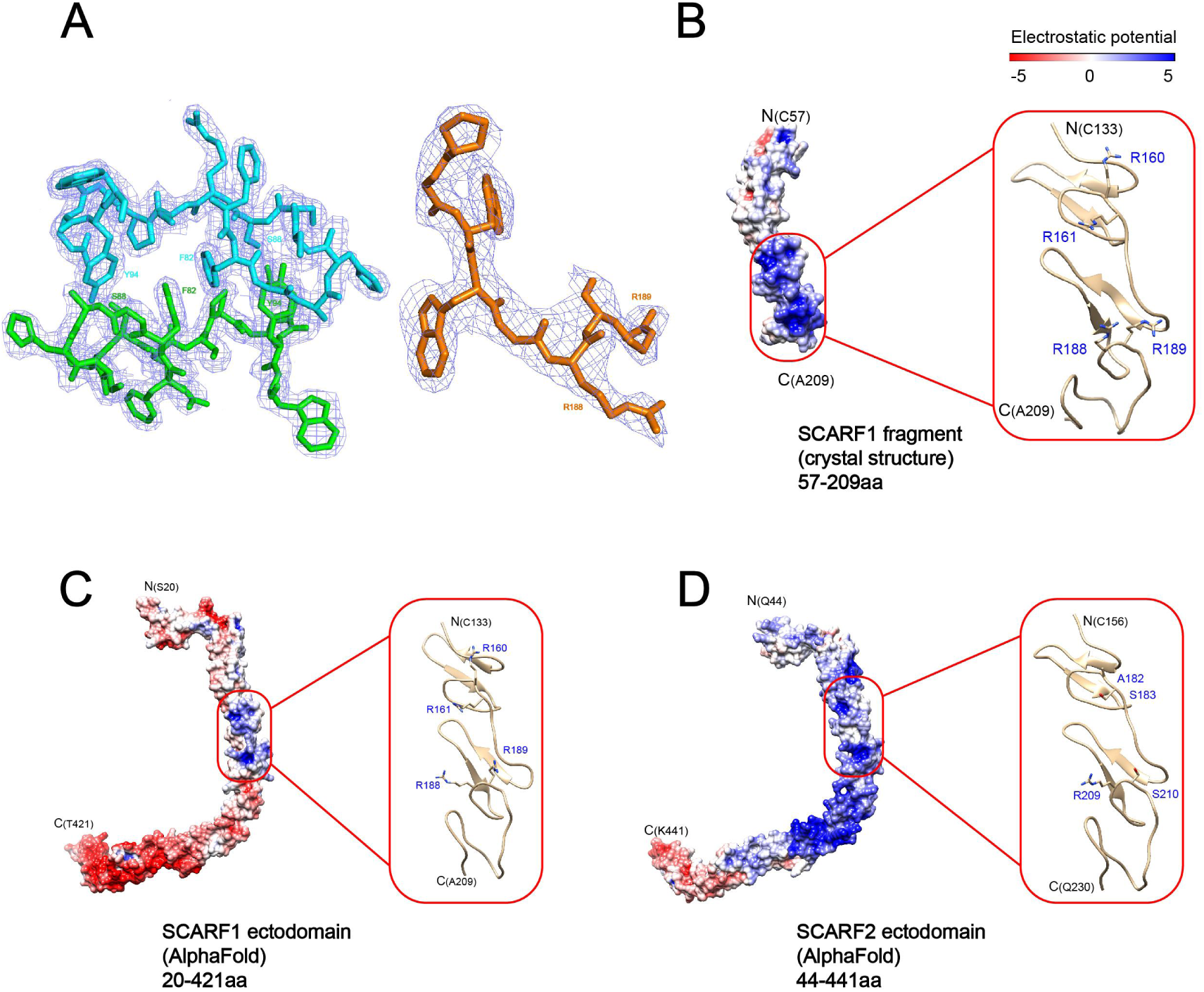
Electron density of the crystals and surface electrostatic potential of SCARF1 and SCARF2. (A) Electron density (light blue) of f1 around the dimeric interface (left) and electron density of f2 at residues (183-189 aa) (right). (B) Surface electrostatic potential of the crystal structure of a SCARF1 fragment (57-209 aa). A ribbon diagram of the region around the lipoprotein binding site (red rectangle) is shown on the right. The side chains of the arginines are also shown. (C) Surface electrostatic potential of the ectodomain of SCARF1 from AlphaFold. A ribbon diagram of the region around the lipoprotein binding site (red rectangle) is shown on the right. The side chains of the arginines are also shown. (D) Surface electrostatic potential of the ectodomain of SCARF2 from AlphaFold. A ribbon diagram of the corresponding region of SCARF1 lipoprotein binding site (red rectangle) is shown on the right. The side chains of the corresponding residues are also shown.

**Figure S3.**
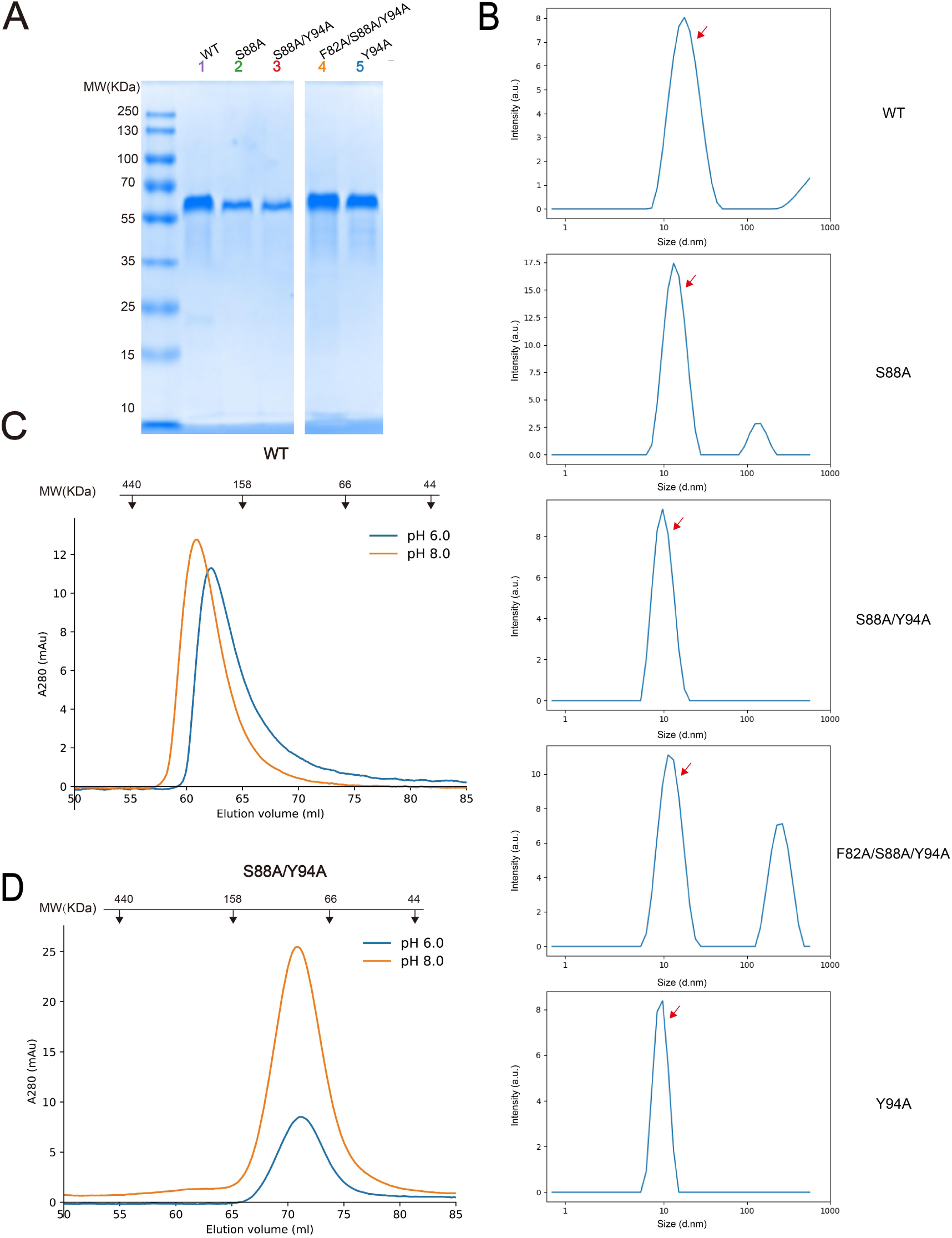
Dimerization of SCARF1. (A) The SDS-PAGE of the peak fractions of SEC profiles of the SCARF1 ectodomain and mutants. (B) The DLS measurements of the wild type and the mutants of SCARF1 ectodomain (red arrows). (C) The SEC profiles of the wild type ectodomain of SCARF1 at pH 6.0 or pH 8.0. (D) The SEC profiles of a monomeric mutant of the ectodomain of SCARF1 at pH 6.0 or pH 8.0. The molecular weight marker of SEC is shown on the top (C-D) (Note: both monomeric and dimeric forms of SCARF1 ectodomain elute earlier than the standard molecular weight marker, probably due to the long conformation of the molecule.).

**Figure S4.**
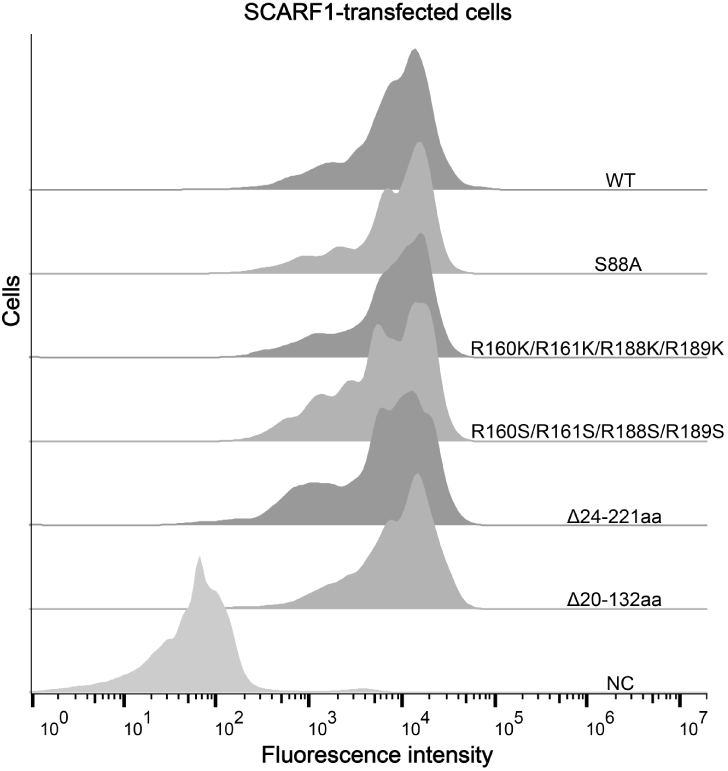
The expressions of SCARF1 and mutants on the cell surface. The expressions of SCARF1 and mutants on the cell surface are monitored by flow cytometry using anti-SCARF1 antibody. The non-transfected cells are included as a negative control (NC).

**Figure S5.**
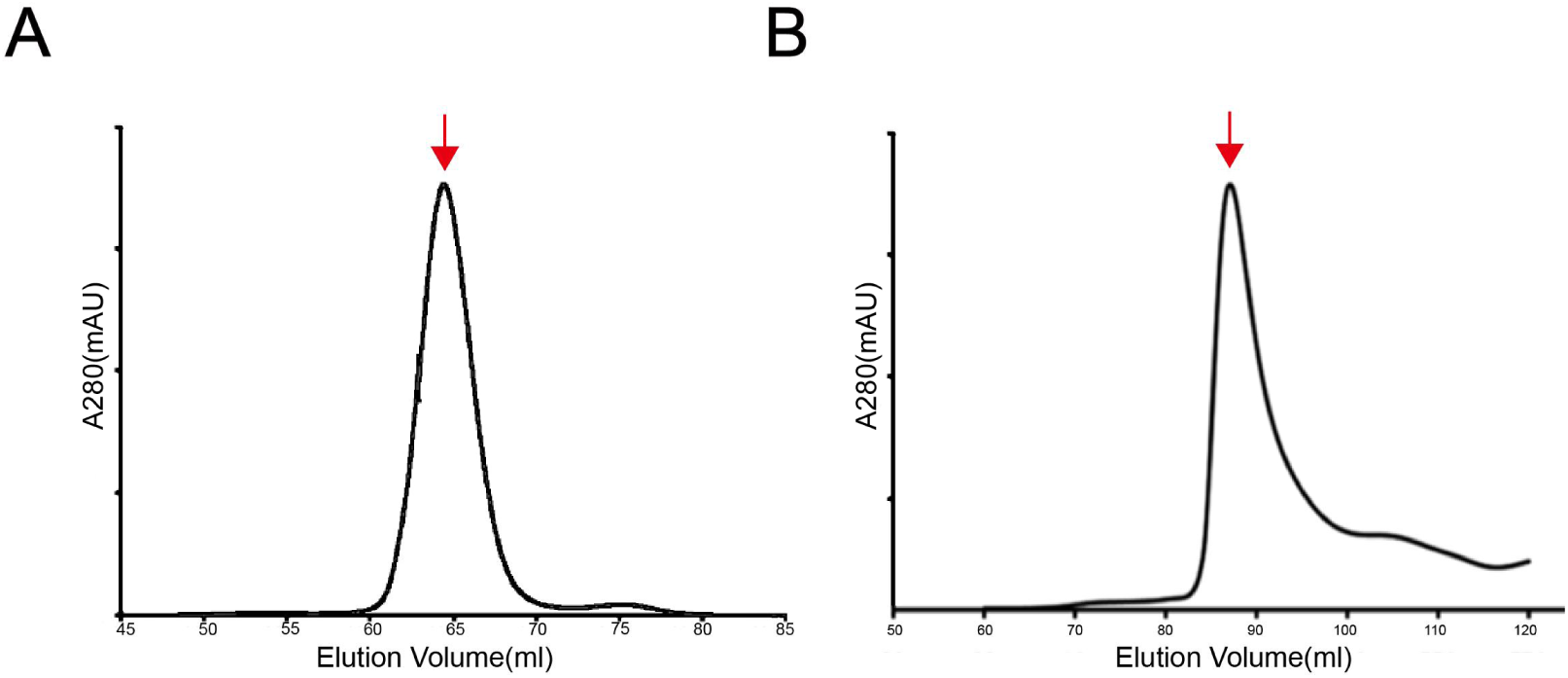
The SEC profiles of the SCARF1 fragments for crystallization. (A) The SEC profile of the purified protein (f1, 20-132 aa) (B) The SEC profile of the purified protein (f2, 20-221 aa)

**Figure S6.**
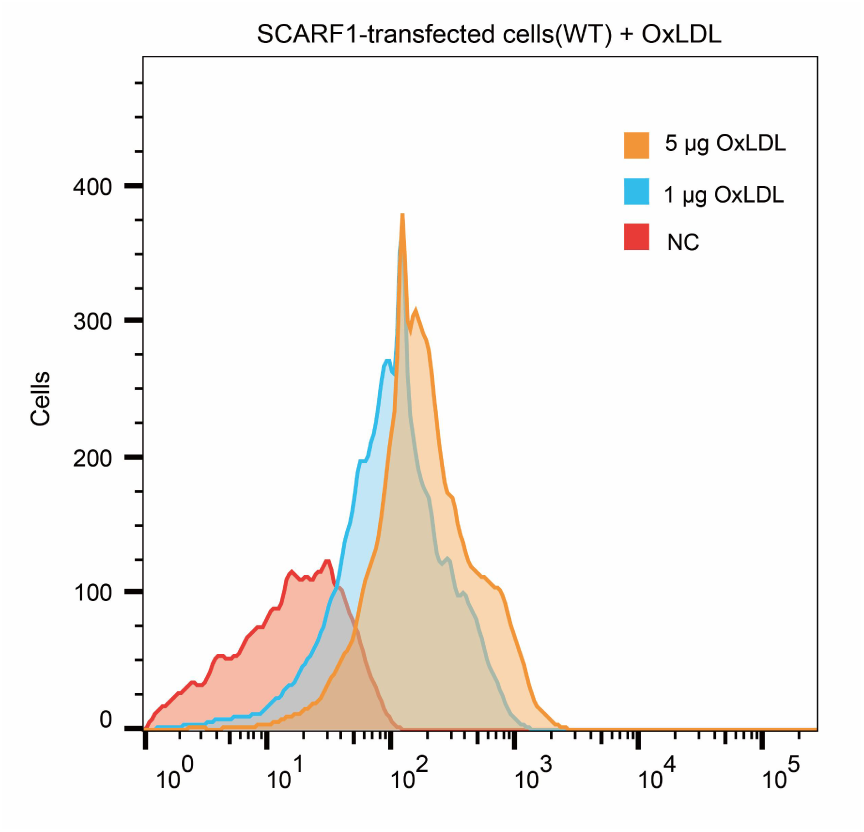
Interaction of OxLDL with the SCARF1-transfected cells by flow cytometry.

**Figure S7.**
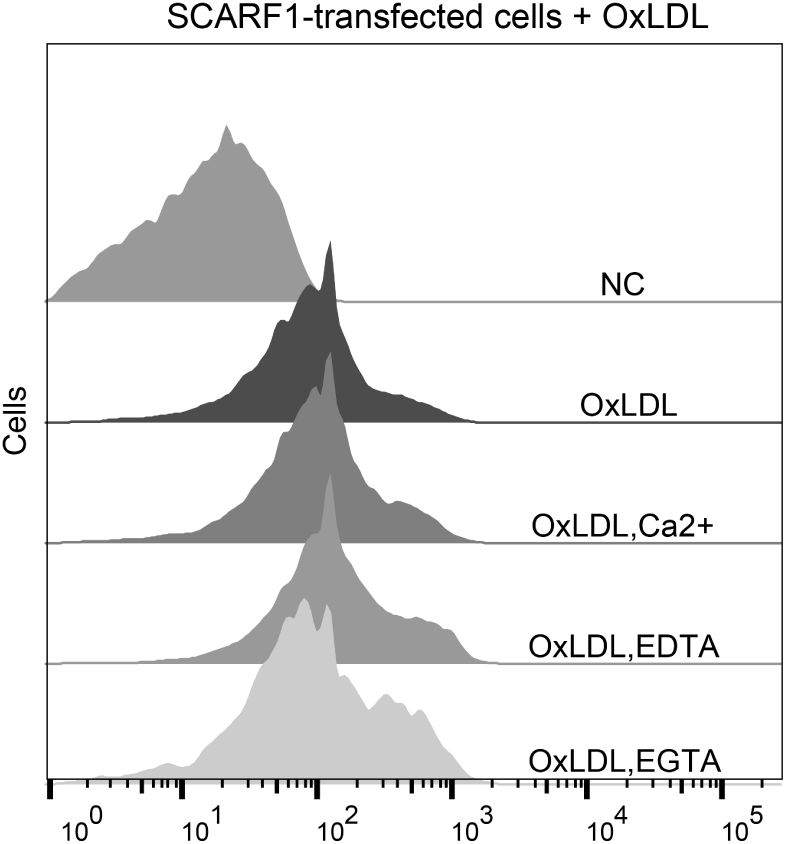
Interactions of OxLDL with the SCARF1-transfected cells in the presence of Ca^2+^, EDTA or EGTA by flow cytometry.

**Table S1.** X-ray data collection and processing.

| Protein | 20-132aa of SCARF1 | 20-221aa of SCARF1 |
| --- | --- | --- |
| Beamline | SSRF BL18U1 | SSRF BL18U1 |
| Wavelength (Å) | 0.98 | 0.98 |
| Space group | P2<sub>1</sub>2<sub>1</sub>2<sub>1</sub> | P4<sub>1</sub>22 |
| Cell parameters |  |  |
| a, b, c (Å) | 47.01, 50.70, 110.74 | 102.01, 102.01, 83.61 |
| $\alpha$, $\beta$, $\gamma$ (°) | 90, 90, 90 | 90, 90, 90 |
| Resolution (Å) | 30.00-2.10<br>(2.18-2.10) | 30.00-2.60<br>(2.69-2.60) |
| R<sub>merge</sub> | 0.075(0.398) | 0.088(0.795) |
| R<sub>pim</sub> | 0.023(0.149) | 0.018(0.176) |
| Unique reflections | 16105(1567) | 14015(1299) |
| I/$\sigma$ (I) | 33(3.03) | 45.5(2.75) |
| Completeness (%) | 99.9(94.6) | 99.2(93.4) |
| Multiplicity | 9.9(7.5) | 23.3(17.9) |

Values in parentheses are for the highest-resolution shells.

**Table S2.** Crystallographic statistics of the structures.

| Protein | 20-132aa of SCARF1 | 20-221aa of SCARF1 |
| --- | --- | --- |
| Resolution (Å) | 24.71-2.20(2.28-2.20) | 26.89-2.60(2.69-2.60) |
| R<sub>work</sub> | 0.219(0.244) | 0.229(0.339) |
| R<sub>free</sub> | 0.252(0.292) | 0.246(0.347) |
| Protein atoms | 3223 | 1133 |
| Wilson B-factor (Å<sup>2</sup>) | 38.8 | 71.2 |
| Average B, all atoms (Å<sup>2</sup>) | 56.0 | 86.0 |
| Rmsd bonds (Å) | 0.36 | 0.43 |
| Rmsd angles (°) | 0.60 | 0.71 |
| Ramachandran favored (%) | 95.41 | 95.36 |
| Ramachandran outliers (%) | 0 | 0 |
| PDB code | 8HN0 | 8HNA |

Values in parentheses are for the highest-resolution shells.

